# Hypernatremia Enhances Transient Outward Potassium and Late Sodium Currents in a Mouse Model of Long QT Syndrome Type 3

**DOI:** 10.64898/2025.12.04.692461

**Authors:** Xiaobo Wu, Sharon A Swanger, Gregory S. Hoeker, Rowan Maisonneuve, Peter Carmeliet, Robert G. Gourdie, Seth H. Weinberg, Steven Poelzing

**Affiliations:** Translational Biology, Medicine, and Health Graduate Program, Virginia Polytechnic Institute and State University, Roanoke, Virginia; Center for Vascular and Heart Research, Fralin Biomedical Research Institute at Virginia Tech Carilion, Roanoke, Virginia, USA; Center for Neurobiology Research, Fralin Biomedical Research Institute at Virginia Tech Carilion, Roanoke, Virginia, USA; Department of Biomedical Sciences and Pathobiology, VA-MD College of Veterinary Medicine, Virginia Tech, Blacksburg, Virginia, USA; Laboratory of Angiogenesis and Vascular Metabolism, Department of Oncology and Leuven Cancer Institute (LKI), KU Leuven, VIB Center for Cancer Biology, VIB, Leuven, 3000, Belgium; Center for Biotechnology, Khalifa University of Science and Technology, Abu Dhabi, United Arab Emirates; Department of Biomedical Engineering and Mechanics, Virginia Polytechnic Institute and State University, Blacksburg, Virginia, USA; Department of Biomedical Engineering, Davis Heart and Lung Research Institute, The Ohio State University, Columbus, Ohio, USA

## Abstract

Cardiac voltage-gated sodium channel gain-of-function (Na_v_GOF) is characterized by action potential duration (APD) prolongation. Hypernatremia and perinexal widening synergistically prolong cardiac APD in guinea pig. However, guinea pig lack the transient outward potassium current (I_to_), which could be increased by hypernatremia and thereby shorten APD.

**Objective:** Determine whether hypernatremia and perinexal expansion synergistically prolong APD in an animal model functionally expressing I_to_.

**Methods:** Whole-cell I_to_ was measured in isolated genetically-modified Na_v_GOF (ΔKPQ) mouse ventricular myocytes. Ventricular APD at 30 (APD30) and 90 (APD90) percent repolarization were measured from optically mapped, Langendorff-perfused wild-type (WT) and ΔKPQ mouse hearts at different perfusate sodium concentrations (145 or 160mM), without and with the perinexal adhesion antagonist βadp1.

**Results:** In isolated myocytes, hypernatremia (160mM sodium) increased I_to_. In whole-heart, hypernatremia significantly decreased both APD30 and APD90 in WT but only APD30 in ΔKPQ preparations. Perinexal disruption with βadp1 did not change APD30 or APD90 in WT hearts, however it decreased APD30 and increased APD90 in ΔKPQ hearts. Combination of hypernatremia and βadp1 did not synergistically change APD in ΔKPQ hearts. Computational models predict that I_to_ activation can prevent synergistic APD prolongation in mouse during hypernatremia and perinexal expansion that was observed previously in a guinea pig Na_v_GOF model lacking I_to_.

**Conclusions:** Hypernatremia during Na_v_GOF prevents early ventricular repolarization due to I_to_ activation (mouse) and prolongs repolarization in the absence of I_to_ (guinea pig). Future studies in animal models electrophysiologically similar to humans are needed to determine if hypernatremia and perinexal expansion are proarrhythmic during Na_v_GOF.

## INTRODUCTION

Cardiac voltage-gated sodium channels are critical for maintaining normal electrical initiation and propagation in hearts.^1^ The sodium channel protein type 5 alpha-subunit Na_v_1.5 is prominently expressed in ventricles,^2^ and mutations associated with this protein are involved in several congenital heart diseases.^3^ In a disease such as congenital Long-QT Type 3 (LQT3), incomplete inactivation of Na_v_1.5 leads to an increased late sodium current (I_NaL_) and constitutes a sodium channel gain-of-function (Na_v_GOF) that prolongs action potential duration (APD) and QT interval. We previously demonstrated in a drug-induced Na_v_GOF guinea pig model that APD can be exacerbated by increasing extracellular sodium concentration and widening the perinexus, a gap junction adjacent extracellular nanodomain in the intercalated disc (ID).^4–6^ In short, when the perinexus widens, sodium ion availability within the cleft increases, and the high density of Na_v_1.5 in the perinexus^7^ allows for an increase in sodium influx (I_NaL_) during incomplete Na_v_1.5 inactivation, resulting in APD prolongation. We also demonstrated that concurrently elevating extracellular sodium and perinexal widening can greatly increase I_NaL_ to synergistically prolong APD.^6^ Therefore, we proposed that managing serum sodium and preventing cardiac edema may prevent arrhythmogenic APD prolongation in patients with congenital and acquired forms of Na_v_GOF.

However, elevating extracellular sodium may concurrently increase the 4-aminopyridine (4-AP) - sensitive transient outward current (I_to_) to shorten APD.^8–11^ Voltage-gated potassium channels such as Kv4.2 and Kv4.3^12^ are responsible for fast and early repolarization after Na_v_1.5 activation by generating I_to_. Importantly, guinea pig myocytes lack classical 4-AP sensitive I_to_, while the current is present in humans.^13–17^ Since increased I_to_ shortens APD, it is unclear if hypernatremia can prolong APD in models of Na_v_GOF where the species functionally expresses I_to_ in the heart.

In the present study, we investigated the individual and combined effects of elevating extracellular sodium and perinexal widening on APD in a genetically-modified Na_v_GOF (ΔKPQ) mouse model.^3^ We confirmed that hypernatremia increases I_to_ in isolated ventricular cardiomyocytes, and this corresponds to hypernatremia-induced APD30 shortening in both WT and ΔKPQ hearts. As a result, hypernatremia *shortens* APD90 in mouse, which is opposite to hypernatremia *prolonging* APD90 in the drug-induced Na_v_GOF guinea pig hearts that lack I_to_.^6^ However, widening the perinexus with the sodium channel β1 subunit de-adhesion peptide (βadp1) prolongs overall APD in ΔKPQ, but not WT hearts, consistent with previous findings in guinea pig. Furthermore, hypernatremia did not shorten APD90 in either WT or ΔKPQ hearts perfused with βadp1, nor did it synergistically prolong APD in the ΔKPQ hearts. The data and computational model suggest that perinexal widening ameliorates APD responsiveness to hypernatremia in a species like mice with a relatively large I_to_. Therefore, we conclude that hypernatremia produces opposing effects on APD in ΔKPQ hearts by increasing I_to_ to shorten APD and increasing I_NaL_ to prolong APD.

## METHODS

The investigation conforms to the Guide for the Care and Use of Laboratory Animals published by the US National Institutes of Health (NIH Publication No. 85–23, revised 1996). All animal study protocols were approved by the Institutional Animal Care and Use Committee at the Virginia Polytechnic Institute and State University.

### Langendorff-perfused heart

Male and female wild-type (WT, SvEv) and ΔKPQ mice^3^ (23-28 weeks of age) were used in this study. Animals were anesthetized with isoflurane inhalation. After cervical dislocation, hearts were rapidly excised, cannulated (<5 minutes) and retrogradely perfused with a crystalloid perfusion solution, consisting of (in mM) 139.5 NaCl, 1.0 MgCl2, 1.2 NaH2PO4, 4.0 KCl, 1.8 CaCl_2_, 10 HEPES; 5.6 Glucose, and 5.5 NaOH (pH=7.4 measured at 36°C). A 145 mM sodium concentration (145Na) is a result of 139.5 mM NaCl and 5.5 mM NaOH, and a high-sodium concentration of 160 mM (160Na) is a result of 154.5 mM NaCl and 5.5 mM NaOH. Importantly, 160Na was chosen to model clinically relevant hypernatremia.^18^ Heart temperature was maintained at 36°C in a three dimensional printed polylactic acid bath.^19^ Atria were removed in order to prevent competitive sinus activation. The perfusion pressure was maintained at 60-80 mmHg by adjusting the flow rate of the perfusion solution. Hearts were stabilized for 15 mins before the experiment started.

### Patch clamp

#### Single cell isolation

Myocytes were enzymatically isolated from the ventricles of adult mouse hearts in Langendorff-perfusion at 37°C with collagenase II. The perfusion buffer for Langendorff-perfused adult mouse hearts consisted of (in mM): 120.3 NaCl, 14.75 KCl, 0.6 KH2PO4, 0.6 Na2HPO4, 1.2MgSO4, 10 HEPES, 4.64 NaHCO3, 30 Taurine, 9.9 2,3-Butanedione 2-monoxime (BDM). The solution pH was adjusted to 7.4 with NaOH. The digestion buffer was made with 135-mg collagenase II (Worthington-biochem, USA) in 60-mL perfusion buffer. Stop buffer was made with 6-mL fetal bovine serum in 54-mL perfusion buffer. A 100-mM CaCl_2_ stock solution was prepared and then 10-μL, 40-μL and 90-μL CaCl_2_ solution were added into three 10-mL stop buffers (CaCl_2_ 100, 400 and 900 μM). Hearts were Langendorff-perfused with the digestion buffer for 8 mins after cannulation. A 3-μL CaCl_2_ stock solution was added in the digestion buffer and then digestion continued for another 8 mins. The digested heart was then placed on a petri dish and gently torn into about 20 pieces. A wide mouth transfer pipette was used to mechanically dissolve the pieces into cells for 2 mins. After 2 mins, a 2-mL stop buffer without CaCl_2_ was added, and the tissue was mechanically agitated for 2 more mins. The cell suspension was transferred to a 50-mL conical tube, and plates were washed with a 5-mL stop buffer. The lysate was spun in conical tubes for 3 mins with a centrifuge at 30G at room temperature. The supernatant was removed and stop buffer added with 100-μM CaCl_2_, and then tubes were re-spun for 3 mins. The last step was repeated with a stop buffer containing 400-μM and then 900-μM CaCl_2_. After removing supernatant, cells were resuspended in 7.5 mL cardiomyocyte media. The media was changed every two hours prior to experiments.

#### Patch clamp protocol

The external solution consisted of (in mM): 1.0 CaCl_2_, 5.5 Glucose, 10 HEPES, 5.4 KCl, 1.0 MgCl2, 145 or 160 NaCl, and 0.33 Na2HPO4. The pH was adjusted to 7.4 with NaOH. Internal solution consisted of (in mM): 1.0 CaCl_2_, 10 EGTA, 5 HEPES, 110 K-Aspartate, 20 KCl, 1.0 MgCl2, 8 NaCl and 5 MgATP. The pH was adjusted to 7.2 with KOH. Myocytes were patch-clamped using the whole-cell configuration in either current-clamp (for action potential recording) or voltage (or action potential)-clamp mode (for I_to_ recordings). To measure I_to_, 3-mM 4-aminopyridine (4-AP) was used to isolate the I_to_ current. The holding potential was set to −80mV for 10 seconds before a 15-ms prepulse to −45 mV in order to inactivate sodium channels and minimize contamination of I_to_.^20^ After 15-ms at −45 mV, voltage was stepped for 500ms between −80 mV and +60 mV in 10 mV increments.

### Optical mapping

The voltage-sensitive fluorophore, di-4-ANEPPS (15 µM, Biotium, CA, USA), was perfused into the heart, and the heart position was visually adjusted to center the anterior epicardial surfaces of both left and right ventricles in the mapping field of view. Action potentials were imaged with epiillumination by a 530nm LED (LEX3G; SciMedia, Costa Mesa, CA) fit with a fiber optic light guide directed through a 520nm bandpass excitation filter, 565nm dichroic excitation mirror, 610nm LP emission filter, and a MiCAM HS02-CMOS camera (SciMedia: 92 x 80 pixels, field of view = 14.02 x 12.19 mm). The electromechanical uncoupler blebbistatin (10 µM, ApexBio Tech LLC, TX, USA), was added to the perfusate to reduce cardiac motion.

In order to investigate epicardial action potential duration (APD) under the same conditions in WT and ΔKPQ hearts, hearts were paced with a unipolar electrode that was placed in the septum near the atrioventricular node to stimulate electrical propagation through the conduction system and the reference electrode was placed in the bath. The stimulation current was set to 1.5X the minimum stimulation threshold which was determined at a cycle length (CL) of 150 ms (pulse width = 2 ms). Each heart was sequentially perfused with different solution compositions. To investigate the effect of hypernatremia on APD, hearts were perfused with 145Na and then 160Na. To investigate APD dependence on extracellular sodium during sodium channel β1-subunit de-adhesion, the peptide βadp1^6,21^ (1 µM, LifeTein LLC, NJ, USA) was perfused to induce perinexal expansion at both sodium concentrations. The perfusion duration of each test solution was 30 mins. Optical signals were recorded during steady-state pacing at CL of 150 ms. For analysis, APD of one optical action potential in each of 1840 pixels was calculated after 2×2 spatial binning by using a custom MATLAB program. APD was measured as the difference between the time of the maximum upstroke velocity (i.e. activation time, AT) and the time to 30% repolarization (APD30) or 90% repolarization (APD90) of the optical action potential (as illustrated in Figure 2A). The average APD across the array was summarized as a single data point for a heart.

### Computational model

A computational murine model simulating WT and Na_v_GOF-associated mutations was used to dissect the roles of hypernatremia on I_NaL_ and I_to_, adapted from our recent study modeling a system of two myocytes, coupled via ephaptic and gap junctional coupling.^22^ Briefly, we accounted for heterogeneous sodium channel subcellular localization by spatially discretizing each cell into an axial membrane patch along the length of the cell and two IDmembrane patches at the ends of the cell. It was assumed that 90% of sodium channels were localized in the ID patches. Extracellular electrical potentials at the ID and cleft are governed by a radial cleft resistance inversely proportional to the intercellular cleft width. ID sodium currents and diffusion with the bulk extracellular space govern the extracellular cleft sodium ion concentration dynamics, which are also modulated by cleft width due to changes in the cleft volume. We utilized a well-established ventricular mouse myocyte ionic model,^23^ representing individual ion channel dynamics, and incorporated a Markov model of a Na_v_GOF mutant (ΔKPQ),^24^ which reproduces a pronounced I_NaL_. Fractional changes in I_to_ are implemented by scaling the I_to_ current conductance. Full model equations, parameters, and simulation codes can be found in our prior work.^22^ The two cells were paced at a CL of 150 ms. Simulations were performed to investigate the relationship of APD with varying sodium concentrations, perinexal (cleft) widths, and fractional I_to_.

### Statistical analysis

Statistical analysis was performed with GraphPad Prism 9 (GraphPad Software Inc., San Diego, CA, USA). The results are presented as mean ± standard deviation. Statistical analyses are described in figure legends. The predominant statistical methods were unpaired Student’s t-tests for comparing between mouse genotypes, paired t-tests for comparing the effect of interventions on hearts, and one-way ANOVA followed by Bonferroni correction for multiple comparisons when analyzing synergistic effects. One-sample *t*-tests were used to determine the individual and combined effects relative to 0. A p<0.05 was considered statistically significant.

## RESULTS

### Effect of Hypernatremia on I_to_ in Isolated Cardiomyocytes

Previous studies demonstrated that hypernatremia can increase the transient outward potassium current (I_to_).^8–11^ To confirm this, I_to_ was measured in isolated ventricular cardiomyocytes obtained from a ΔKPQ heart exposed to 145 and 160 mM extracellular sodium concentrations. Current-voltage (I-V) curves of I_to_ under conditions of 145Na and 160Na are shown in Figure 1 (left). 160Na shifted the I-V curve upward suggesting that hypernatremia increases I_to_. Furthermore, I_to_ at +60 mV was significantly higher with 160Na than 145Na (Figure 1– right). Overall, we confirm that hypernatremia increases I_to_ in isolated mouse cardiomyocytes.

**Figure 1.**
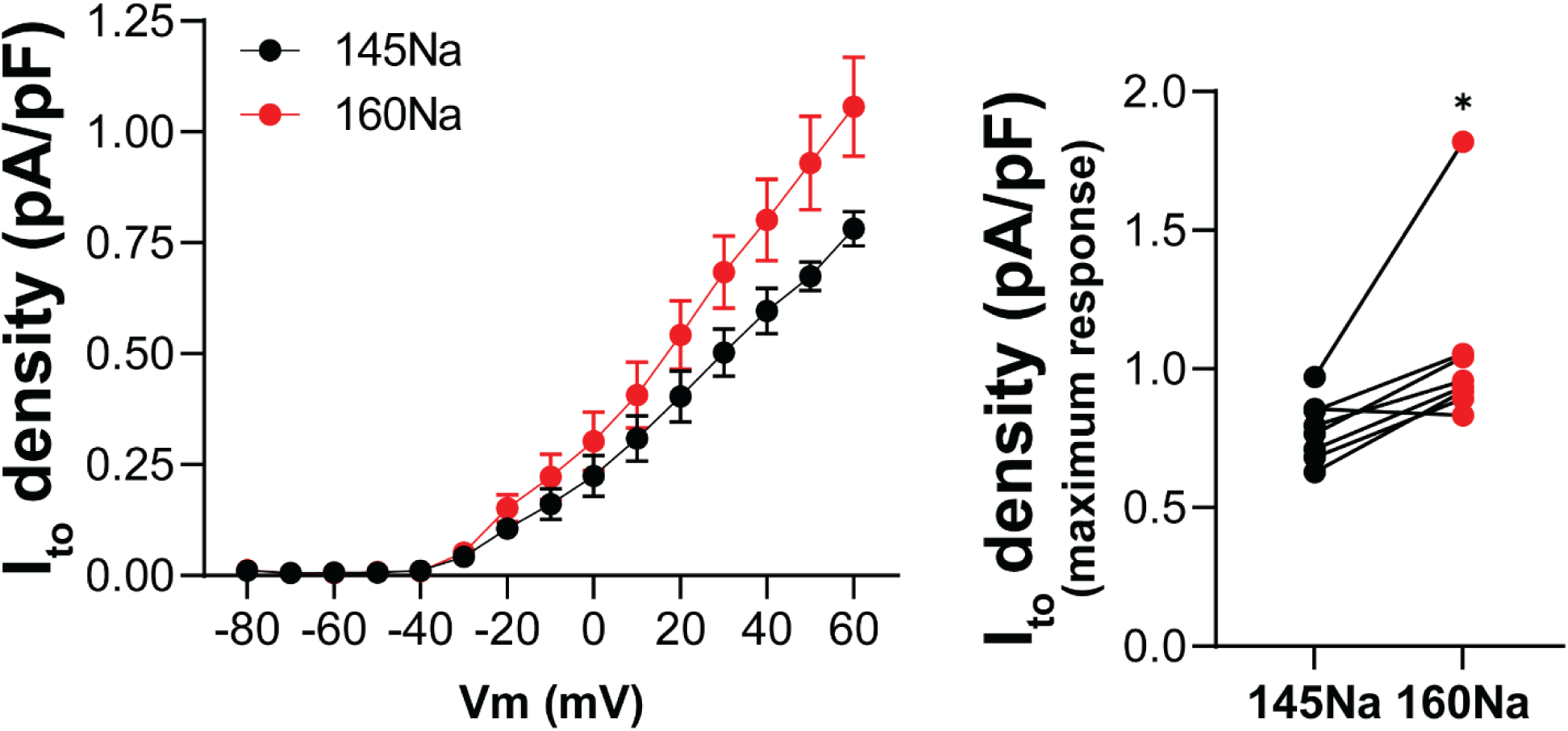
Hypernatremia increases the transient outward potassium current (I_to_). **Left:** I-V curves of I_to_ under conditions of different sodium concentrations from isolated ventricular cardiomyocytes (n=8) of a ΔKPQ heart. **Right:** I_to_ at +60 mV was significantly increased by hypernatremia (160Na). *p<0.05 (paired Student’s *t* test).

### Effect of Hypernatremia on APD in Whole-Heart Preparations

Epicardial action potentials from anterior left and right ventricles were obtained by optically mapping Langendorff-perfused WT and ΔKPQ mouse hearts. Representative action potential tracings and APD contour maps (APD30 and APD90) from WT and ΔKPQ hearts are shown in Figure 2A and 2B, respectively. Previous studies reported I_to_ differs by sex in mouse ventricular myocytes, with males exhibiting larger I_to_ than females.^25,26^ Based on this, we analyzed APD separately by sex in both WT and ΔKPQ hearts. Interestingly, our data did not reveal any significant sex-related APD differences in either genotype (Figure S1), and therefore, data are pooled. Overall, comparison of APD30 and APD90 from all WT and ΔKPQ hearts at a baseline sodium concentration of 145 mM demonstrated that APD is significantly longer in ΔKPQ hearts relative to WT (Figure 2C).

**Figure 2.**
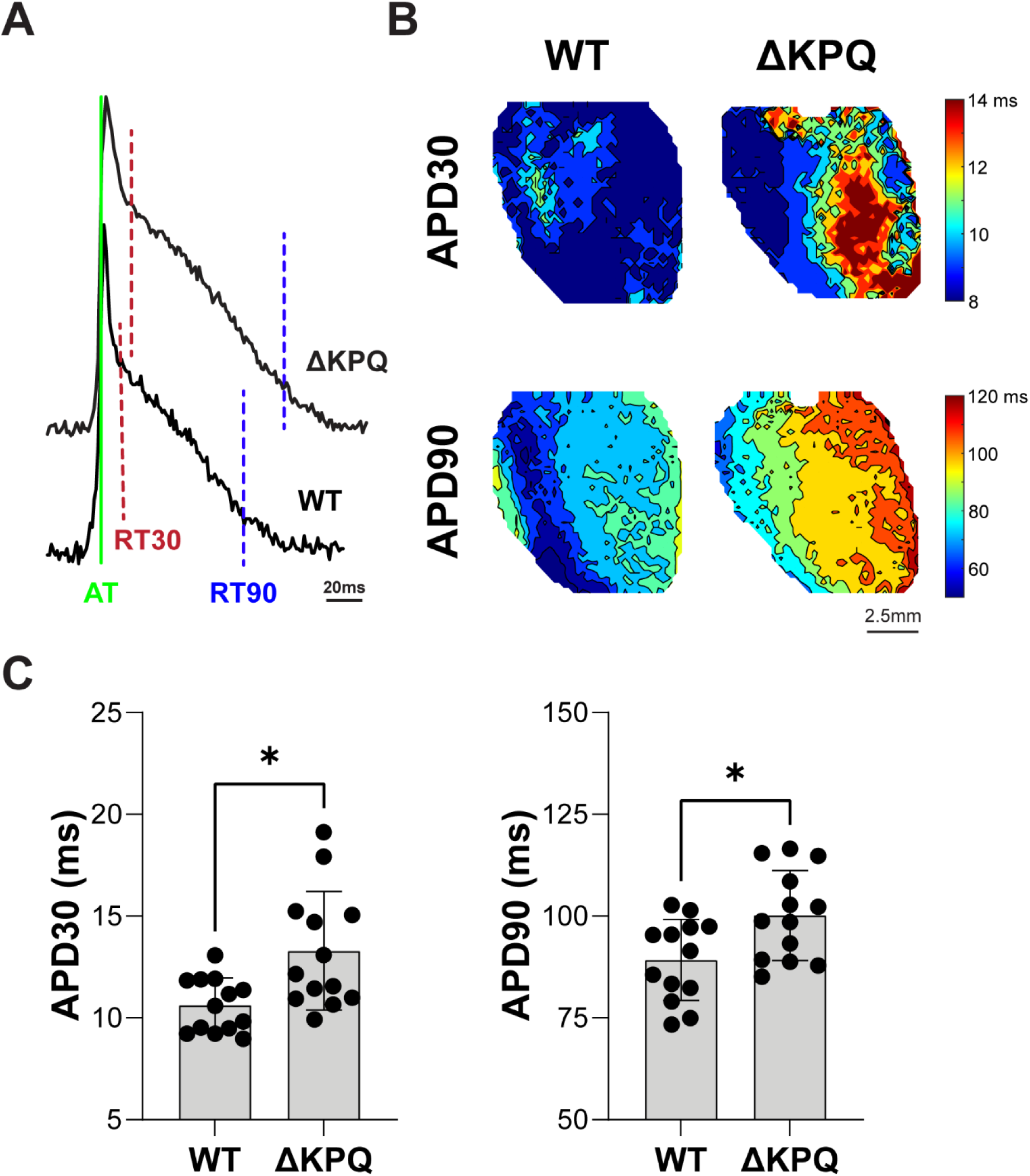
The phenotype of prolonged APD is manifested in ΔKPQ mouse hearts. **A.** Ventricular action potential duration (APD) was quantified with APD30 (the difference between activation time -AT and 30% repolarization time -RT30) and APD90 (the difference between AT and 90% repolarization time -RT90) in WT and ΔKPQ hearts. **B.** Representative ventricular APD30 and APD90 contour maps from a WT and a ΔKPQ heart. **C.** APD30 and APD90 were significantly greater in ΔKPQ (n=13) than WT hearts (n=13). *p<0.05 (unpaired Student’s *t* test). AT, activation time; RT30 and RT90, 30% and 90% repolarization time.

To determine if hypernatremia either shortens APD due to I_to_ activation or prolongs APD via an ID mechanism as suggested by Na_v_GOF guinea pig studies and models,^4–6^ extracellular sodium was raised to 160 mM (160Na). Consistent with the hypernatremic enhancement of I_to_, 160Na decreased APD30 in both WT and ΔKPQ hearts relative to 145 mM sodium (145Na) (Figure 3A and 3B). 160Na also significantly decreased APD90 in WT hearts. By contrast, hypernatremia alone did not significantly change APD90 in ΔKPQ hearts. In summary, I_to_ enhancement by hypernatremia facilitates early (shorter APD30) and late repolarization (shorter APD90) in WT animals. In the ΔKPQ hearts, hypernatremic I_to_ activation only enhances early repolarization. Thus, despite what appears to be species specific differences in response to hypernatremia during Na_v_GOF, the data are consistent that hypernatremia during Na_v_GOF can facilitate early ventricular repolarization in a species functionally expressing I_to_ (mouse) or prolong it without functional I_to_ expression (guinea pig).^6^

**Figure 3.**
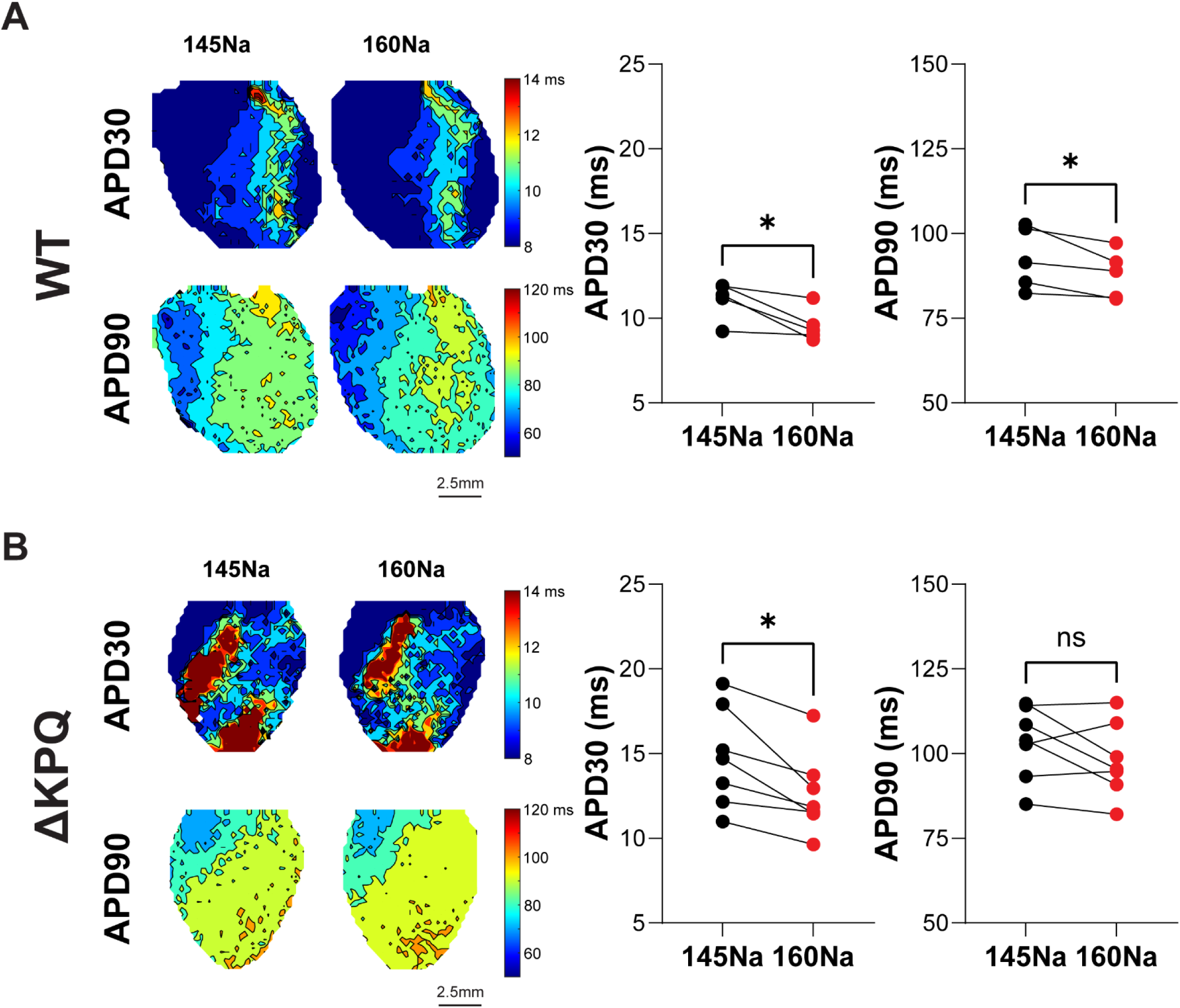
Increased extracellular sodium shortens APD differently in WT and ΔKPQ mouse hearts. **A-Left:** Representative contour maps of ventricular APD30 and APD90 from a WT heart at baseline (145Na) and during hypernatremia (160Na). **Right**: Hypernatremia significantly decreased APD30 and APD90 in WT hearts (n=5). **B-Left:** Representative contour maps of ventricular APD30 and APD90 from a ΔKPQ heart at baseline (145Na) and during hypernatremic conditions (160Na). **Right:** Hypernatremia significantly decreased APD30 but not APD90 in ΔKPQ hearts (n=7). 145Na and 160Na, 145 mM and 160 mM sodium concentrations, respectively. *p<0.05 (paired Student’s *t* test). ns, not significant.

### Effect of Perinexal Expansion on APD in Whole-Heart Preparations

We previously demonstrated that widening the perinexus prolongs APD in the guinea pig drug-induced Na_v_GOF model by a mechanism of attenuating sodium ion depletion from the ID, which permits enhanced I_NaL_.^4,6,27^ Furthermore, widening the perinexal cleft and increasing extracellular sodium “synergistically” increased APD more than the sum of the individual effects. In summary, sodium ion availability in diffusion limited extracellular clefts can be increased by increasing extracellular sodium concentration, widening the perinexus, or both.

Under baseline conditions (145Na), we found that the perinexal widening peptide βadp1 did not change APD30 or APD90 in WT hearts (Figure 4A), which is consistent with previous findings.^6,21^ By contrast, βadp1 decreased APD30 while concurrently increasing APD90 in ΔKPQ hearts (Figure 4B). Thus, βadp1 prolonged final repolarization measured by APD90 in ΔKPQ hearts, also consistent with previous findings.^6^

**Figure 4.**
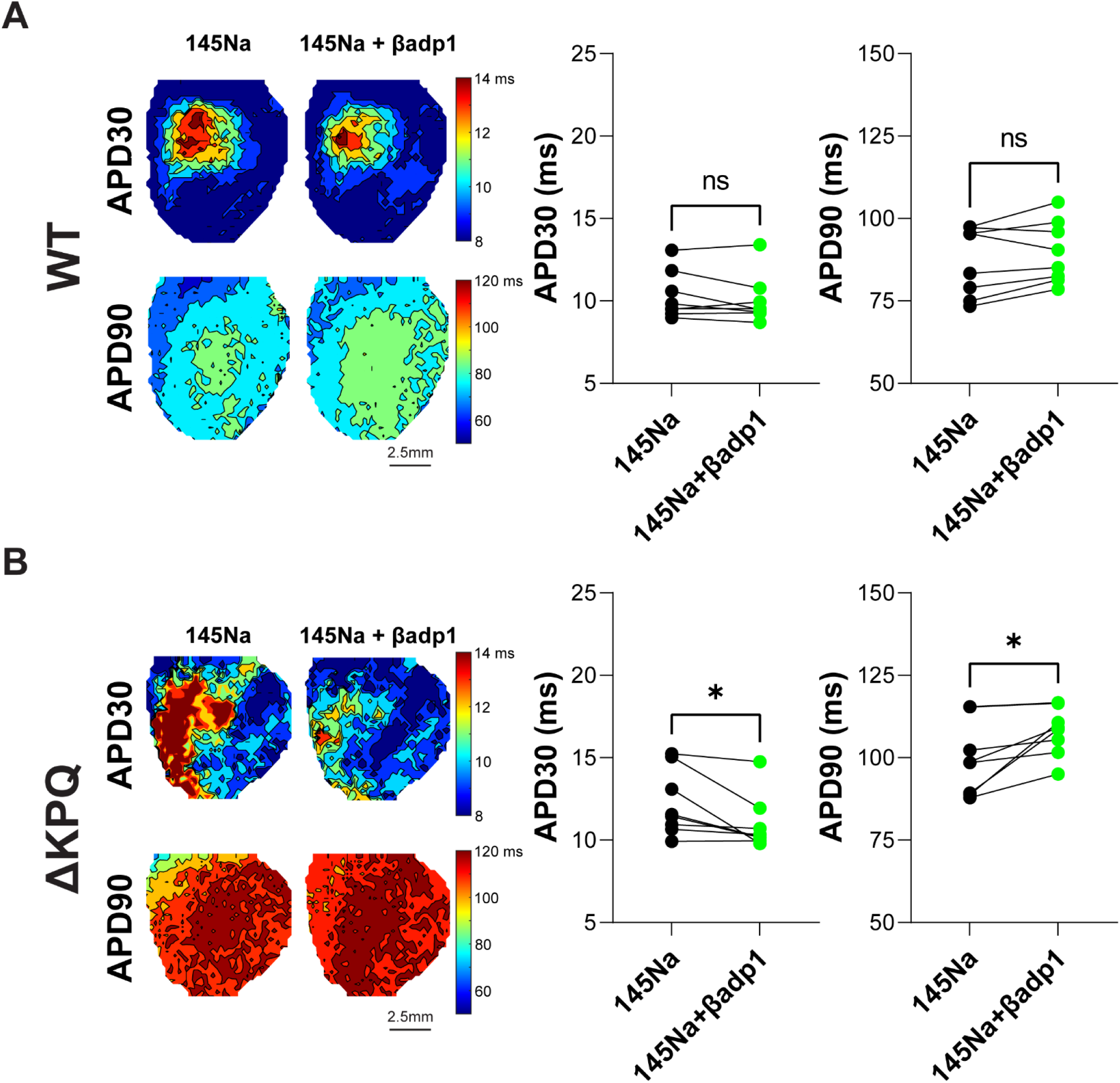
Application of βadp1 changes APD in ΔKPQ but not WT mouse hearts. **A-Left:** Representative contour maps of ventricular APD30 and APD90 from a WT heart at baseline (145Na) and during treatment with 1 µM βadp1. **Right:** βadp1 had no significant effect on APD30 or APD90 in WT hearts (n=8). **B-Left:** Representative contour maps of ventricular APD30 and APD90 from a ΔKPQ heart at baseline (145Na) and during treatment with 1 µM βadp1. **Right:** βadp1 significantly decreased APD30 and increased APD90 in ΔKPQ hearts (n=8). *p<0.05 (paired Student’s *t* test). ns, not significant.

### Effect of Combined Hypernatremia and Perinexal Expansion on APD in Whole-Heart Preparations

Next, hypernatremia and βadp1 were combined in this murine model to determine whether these interventions synergistically prolong APD, as observed in the guinea pig drug-induced Na_v_GOF model.^6^ Figure 5 summarizes the change in APD relative to hearts perfused at baseline with 145Na. In WT hearts, elevating sodium to 160Na decreases APD30 and APD90 more than perfusing with βadp1 at 145Na (Figure 5A). Second, the decrease in APD30 when sodium was increased in WT heart perfused with βadp1 (160Na+βadp1) was significantly different from zero, but not significantly different from the effect of βadp1 alone. Furthermore, the combination of elevating sodium and perinexal expansion (160Na+ βadp1) did not change APD90 relative to either elevating sodium (160Na) or perfusing with βadp1 alone.

**Figure 5.**
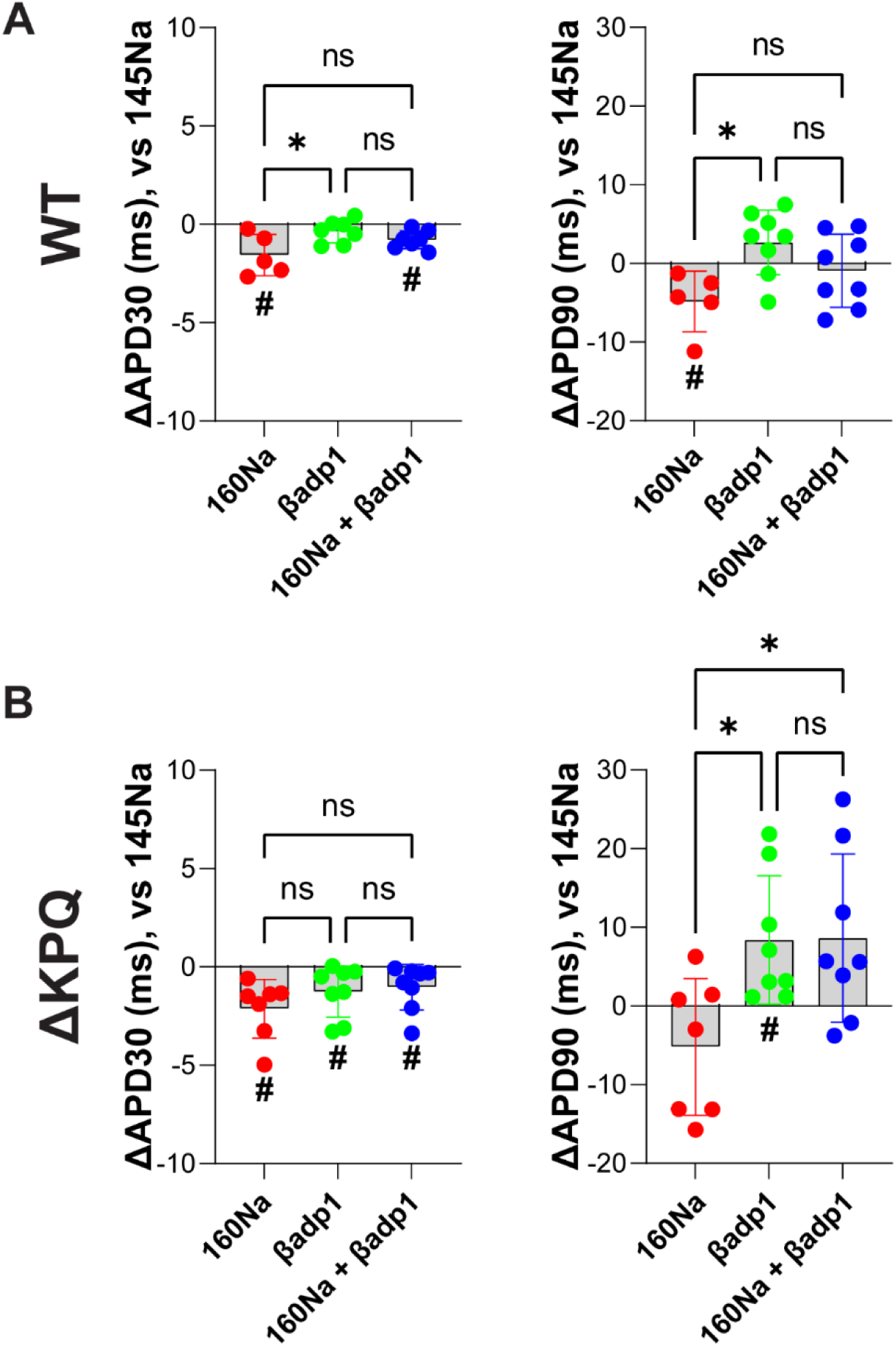
Combination of hypernatremia and βadp1 has no synergistic effect on APD in ΔKPQ mouse hearts. **A:** Combination of hypernatremia (160Na) and βadp1 (1 µM, n=7) decreased APD30 but not APD90 in WT hearts**. B:** Combination of hypernatremia (160Na) and βadp1 (1 µM, n=8) decreased APD30 and did not significantly increase APD90 in ΔKPQ hearts, although the change in APD90 was more positive when comparing 160Na+βadp1 to 160Na. *p<0.05 (One-way ANOVA with Bonferroni correction for multiple comparisons). #p<0.05 (One sample *t* test relative to 0). All changes in APD30 and APD90 relative to baseline values at 145Na.

In the ΔKPQ hearts, raising extracellular sodium similarly reduced APD30 as did perfusion of βadp1 alone or perfusion of βαdp1 with 160Na relative to hearts perfused with 145Na alone (Figure 5B). Interestingly, APD30 and APD90 in ΔKPQ hearts responded differently to βadp1. Specifically, while the degree of APD90 increase with 160Na+βadp1 relative to 145Na was not significantly different from zero, it was significantly more positive than the effect of increasing extracellular sodium alone (160Na). Taken together, these data suggest that perinexal expansion is sufficient to marginally, but not significantly, attenuate APD90 sensitivity to hypernatremia in WT animals, and to statistically counteract the APD90 shortening effect of I_to_ enhancement by hypernatremia in a mouse model of Na_v_GOF.

### Computational Modeling Predictions

At present, the response of APD to hypernatremia and perinexal expansion in a murine model with I_to_ is different from that obtained in guinea pig lacking I_to_. To predict how I_to_ enhancement by hypernatremia modulates APD in murine ventricular myocardium, a computational model simulating a 1-dimensional strand of coupled myocytes was employed. Figure 6 demonstrates that enhancing late sodium current (WT compared with ΔKPQ; dashed to solid lines) increases APD30-90 for all perinexal widths, similar to previously published guinea pig models^4,6,28^ and human data.^29^ Increasing extracellular sodium in WT (Figure 6, black to red dashed lines) negligibly decreases all APD values for all perinexal widths. In the ΔKPQ myocytes, increasing sodium (black to red solid lines) also modestly decreases APD30 for all perinexal widths, similar to WT (dashed). However, increasing sodium (black to red) fundamentally alters the APD50-90 dependence on perinexal width in ΔKPQ myocytes (solid), relative to WT (dashed) myocytes, such that hypernatremia generally increases APD50, APD70, and APD90 for all but the narrowest perinexi. While increasing extracellular sodium (black to red) can increase I_NaL_ by well-established mechanisms of increasing the driving force for sodium, increasing perinexal width opposes this effect by enhancing repolarization. It does so by preventing the rapid depletion of cleft sodium concentration that occurs in diffusion limited clefts consequent to sodium withdrawal into the intracellular space by I_NaL_, thereby permitting a sustained I_NaL_, as has been previously described.^4^

**Figure 6.**
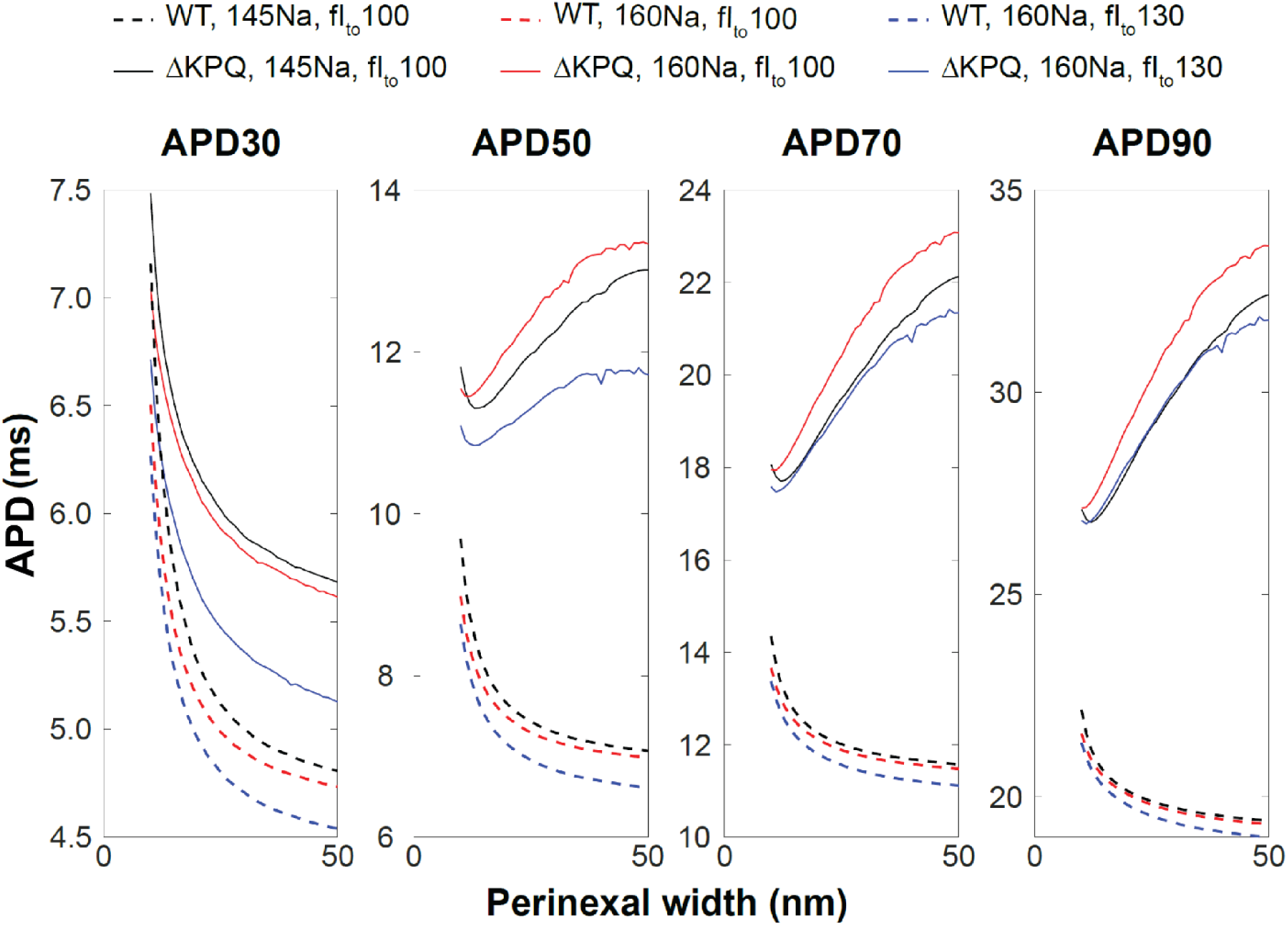
Enhanced I_to_ attenuates APD sensitivity to hypernatremia during sodium channel gain-of-function in a mouse ventricular action potential model. Enhancing late sodium current (dashed to solid lines) increases APD30-90 for all perinexal widths. Increasing extracellular sodium in WT (black to red) modestly decreases all APD values in WT (dashed) for all perinexal widths. In the ΔKPQ myocytes, increasing sodium (black to red) decreases APD30 for all perinexal widths, while APD50-90 increase. Increasing the fractional conductance of I_to_ (fI_to_ - red to blue) decreases APD30-90 and abolishes or even reverses APD prolongation caused by increasing extracellular sodium.

Note the simulations described so far (black, red lines) do not incorporate any potential influence of hypernatremia on I_to_. Increasing the fractional conductance of I_to_ in the ΔKPQ simulated myocytes from 100 (baseline) to 130% (enhanced; fI_to_ – solid red to blue) decreases APD30-90 without fundamentally altering APD dependence on perinexal width. Importantly, increasing I_to_ not only abolishes APD prolongation for different stages of repolarization, it can even modestly reverse APD90 prolongation caused by increasing extracellular sodium during Na_v_GOF. That is, simulations demonstrate that the combination of enhancement of I_NaL_ by hypernatremia (due to increased driving force) and I_to_ (via increased conductance) result in ΔKPQ mouse myocytes with near identical APD90 values for all perinexal widths (c.f., black and blue solid lines). In summary, simulations predict that enhancing I_to_ by 30%, comparable to the average increase measured in Figure 1 (145Na vs 160Na: 0.78±0.05 vs 1.05±0.14 pA/pF at 60mV), is sufficient to attenuate APD sensitivity to hypernatremia and perinexal expansion during Na_v_GOF in a mouse ventricular action potential model.

Examination of the action potential, I_Na_, and I_to_ illustrate the mechanisms underlying the changes in APD during hypernatremia and enhanced I_to_ (Figure 7). In ΔKPQ myocytes (solid lines), increasing extracellular sodium (red and blue lines) produces a modest increase in I_Na_ due to larger sodium driving force, which leads to an early action potential peak, that in turn enhances I_to_ (Figure 7C), thereby shortening APD30 (solid red, note overlap of blue and red solid traces during the upstroke in panels A and B). As noted above, increasing extracellular sodium also enhanced late I_Na_ (Figure 7B, inset), such that APD90 is comparable between baseline (black) and hypernatremia with enhanced I_to_ (blue).

**Figure 7.**
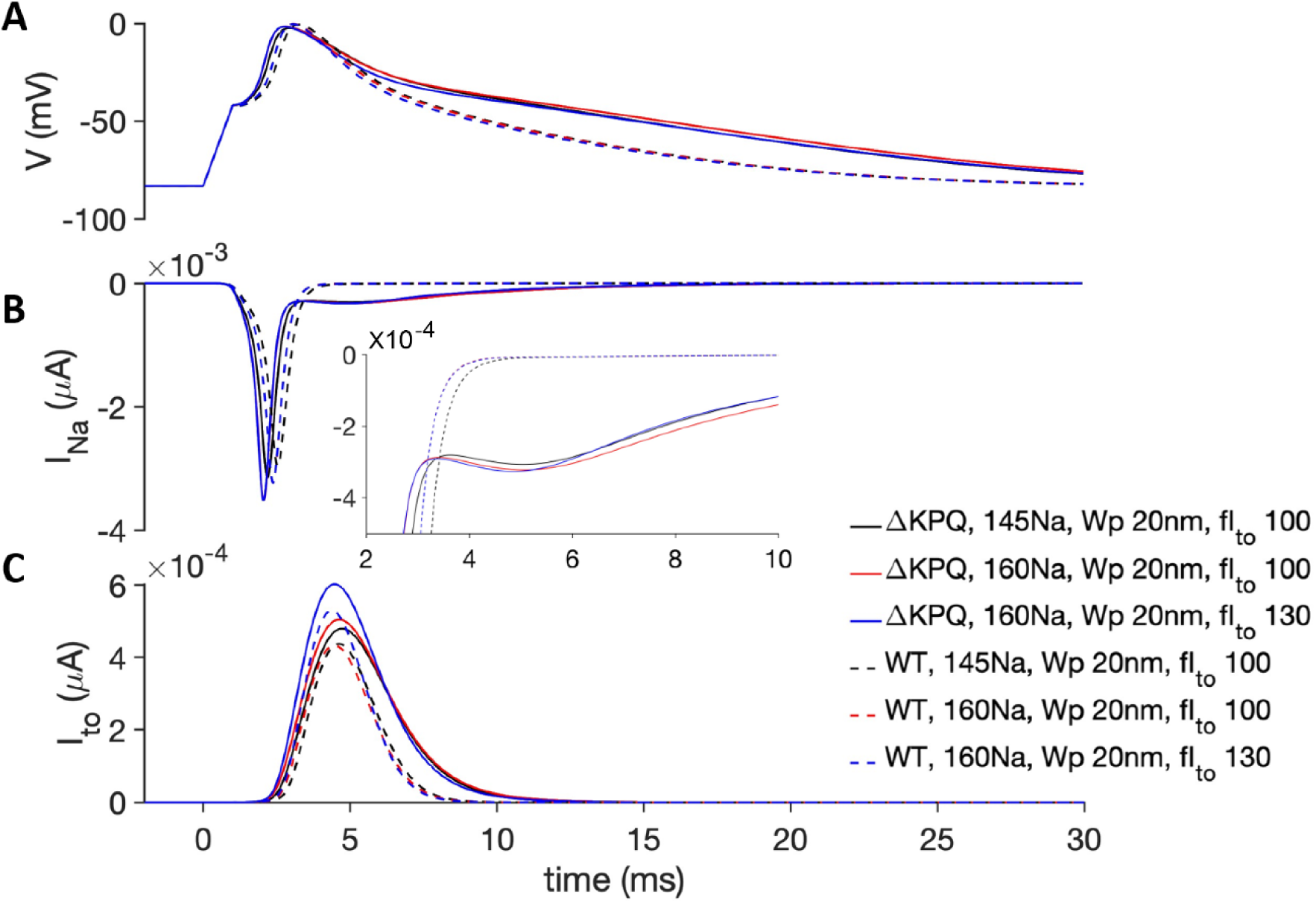
Hypernatremia enhanced late I_NaL_ and I_to_ during sodium channel gain-of-function in a mouse ventricular action potential model. (A) Action potential, (B) I_Na_, and (C) I_to_ are shown for ΔKPQ (solid) and WT (dashed) for baseline (black), hypernatremic (red, blue), and enhanced I_to_ (blue) conditions. Note red and blue solid lines mostly overlap in panels A and B during the action potential upstroke.

## DISCUSSION

We previously demonstrated that elevating extracellular sodium with and without perinexal widening prolongs action potential duration (APD) in drug-induced sodium channel gain-of-function (Na_v_GOF) guinea pig hearts,^6^ a species which does not functionally express transient outward potassium channels.^13–17^ By contrast, we found in the present study that hypernatremia shortened APD in WT but only early repolarization in ΔKPQ mouse hearts, and this is likely due to enhanced transient outward potassium current (I_to_). Furthermore, the synergistic APD prolonging effect of hypernatremia and βadp1 found in the drug-induced Na_v_GOF guinea pig model was not evident in ΔKPQ mice. We attribute this species difference to the absence of the 4-AP sensitive I_to_ channels in guinea pig and substantial expression of I_to_ channels in mice.

The mechanism by which increasing sodium ion concentration can enhance I_to_ is still an active area of investigation.^8–11,30,31^ One hypothesis of confounding relevance is associated with enhanced extracellular sodium leading to intracellular calcium depletion and subsequent post-translational modification of I_to_ through calcium sensitive kinases such as the calcium/calmodulin-dependent protein kinase II (CaMKII). If elevated extracellular sodium indirectly decreases CaMKII via enhanced forward mode function of the sodium-calcium exchanger, this should decrease I_to_ peak current and facilitate I_to_ inactivation as previously demonstrated.^32,33^ Yet, we report for the first time in mice that increasing extracellular sodium increases I_to_ as previously reported in other species.^8–11^ While the functional implications of CaMKII are not the major focus of this study, the discussion of this point is to illustrate that intracellular calcium remodeling induced by hypernatremia may have effects on cardiac electrophysiology beyond modulating I_to_.

We also demonstrated that widening the perinexus with βadp1 did not significantly change APD in WT while increasing APD90 in ΔKPQ hearts. This finding is consistent with previous studies in drug-induced Na_v_GOF where perinexal expansion by similar and different experimental interventions exacerbates APD prolongation.^6,21^ Interestingly, we found that βadp1 decreased APD30 in ΔKPQ hearts, which could also be caused by increased I_to_. Specifically, the βadp1 peptide was designed to interfere with the sodium channel beta1 subunit Navβ1 encoded by SCN1B. Importantly, Deschenes et al demonstrated that Navβ1 increases peak I_to_, and silencing Navβ1 reduces functional expression of Kv4.2, Kv4.3 and KChIP2 mRNA and associated proteins.^34,35^ Thus, if βadp1 interferes with Scn1b associated with Na_v_1.5, it is possible that it also interacts with I_to_ to increase current density. However, this effect was not observed in our WT hearts. Specifically, neither APD30 nor APD90 were changed by βadp1 in WT hearts. At present, it remains unknown why βadp1 does not significantly decrease APD30 in WT but shortens APD30 in ΔKPQ. Furthermore, although APD30 decreases with βadp1, APD90 still prolongs in the ΔKPQ mouse heart with perinexal expansion, suggesting that final repolarization in the Na_v_GOF mouse model is mainly mediated by enhanced I_NaL_ caused by perinexal widening.

In the absence of tools to directly measure transmembrane currents and extracellular ionic concentrations in nanodomains as narrow as 20 nm between myocytes, we turned to computational modeling to explore the dynamics of electrical potentials and currents in these nanodomains and offer insights into likely mechanisms underlying experimental results. In our mouse computational model that includes ephaptic coupling and an extracellular cleft space, we demonstrate that increasing extracellular sodium can shorten APD30 without changing APD90 due to enhancement of both I_NaL_ and I_to_. As has been demonstrated previously, enhancing the action potential phase 1 notch due to I_to_ can increase as well as decrease action potential duration, depending on I_to_ level.^36^ Regardless, the models further predict that as little as a 30% increase in I_to_ current density can abolish APD prolongation by enhanced I_NaL_ and wide ID clefts. In other words, hypernatremia increases I_NaL_ at IDs, but also enhances outward I_to_ in murine hearts such that the hypernatremia and wide perinexi synergy that is observed in guinea pig is absent in mice (Figure 8).

**Figure 8.**
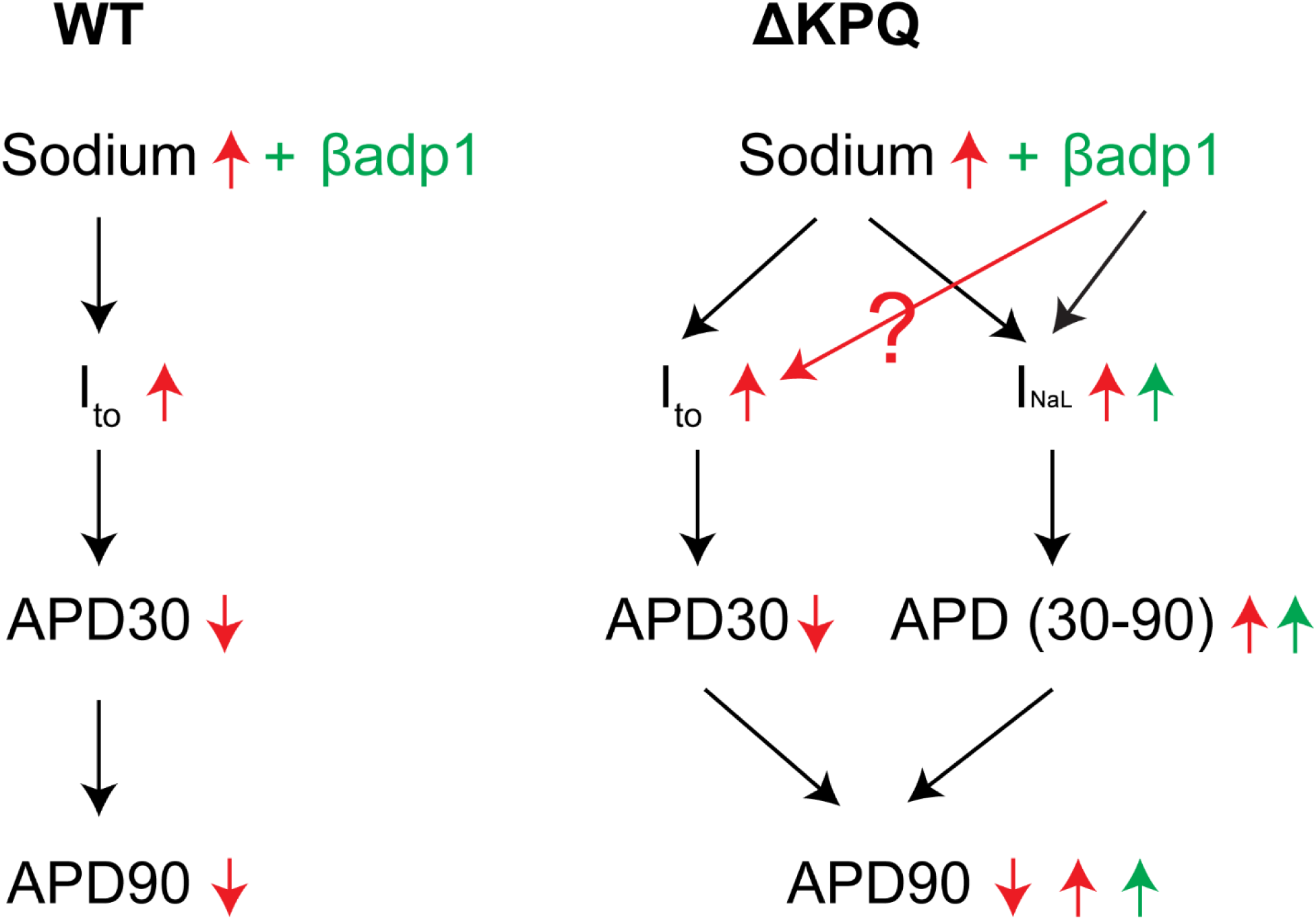
The proposed mechanism of the individual and combined effects of hypernatremia and βadp1 on APD in WT and ΔKPQ hearts. Red arrows indicate the effect of high sodium; Green arrows indicate the effect of βadp1; Question mark indicates unclear effect of βadp1 on I_to_.

The previously reported synergistic effects of hypernatremia and βadp1 on Na_v_GOF phenotype led to speculation that modulating plasma sodium in patients with congenital and acquired forms of Na_v_GOF may be therapeutic. However, this was based on a guinea pig model of Na_v_GOF, and guinea pig ventricular myocardium does not functionally express I_to_, as mentioned earlier. It is therefore important to understand the electrophysiological differences between species to account for the different effects of hypernatremia on APD during Na_v_GOF. First, while humans functionally express I_to_, murine myocardium expresses significantly more I_to_ relative to other animals.^37^ As a result, I_to_ is primarily responsible for the early repolarization and “triangular” action potential in murine cardiomyocytes. Second, there is no action potential plateau in mouse cardiomyocytes relative to other animals such as guinea pig, rabbit, and human. Simulations predict that the action potential plateau is critical for maintaining I_NaL_ during repolarization in order to prolong APD. Early repolarization can shift sodium channels from inactivated to deactivated states to close the channel and reduce I_NaL_. Additionally, we previously demonstrated that Na_v_1.5 activation during the upstroke of the action potential can rapidly deplete sodium ions in narrow ID nanodomains like the perinexus, via a process called ephaptic self-attenuation.^4^ This self-attenuation could limit I_NaL_ and thereby also prevent APD prolongation during a longer action potential. This mechanism may be enhanced in mice which lack a long action potential and the ability for sodium to refill in the cleft. Therefore, the unique differences in cardiac electrophysiology between mouse and other animal hearts suggest that the translational potential of some experiments are model-dependent. Taken together, data from the present study conducted with mice and previous studies on guinea pig Na_v_GOF suggest that extracellular sodium may be an arrhythmic risk factor in diseases also associated with I_to_ loss of function as occurs in heart failure, or gain-of-function as can occur in the Brugada Syndrome.^38^ Importantly, revealing the effects of extracellular sodium on the manifestation of an arrhythmogenic repolarization substrate in humans is necessary to determine whether plasma ionic composition is an important biomarker for predicting arrhythmia incidences associated with sodium and transient-outward potassium channel remodeling or channelopathies.

There are several limitations in the current study. First, measurement of perinexal width was not conducted as previous studies have demonstrated perinexal expansion in guinea pig and mice associated with the osmotic agents, βadp1, and genetic Scn1b ablation.^5,6,21,39–41^ Furthermore, the finding that βadp1 had no effect on APD in WT mice, while APD90 prolonged in ΔKPQ mouse hearts, is consistent with the same effect observed with perinexal expansion in drug-induced guinea pig Na_v_GOF studies, suggesting that βadp1 widens the perinexus. Second, we did not investigate the mechanism by which hypernatremia increases I_to_ in WT mice hearts experimentally, but the effect of I_to_ enhancement by hypernatremia has been observed in other species as mentioned previously. ^8–11^ Third, we did not investigate how other ionic currents such as L-type calcium channels or potassium channels are modulated by hypernatremia. Lastly, since I_to_ current density is larger in mice relative to humans, future studies in humans are necessary to determine correlation between acute APD remodeling and plasma sodium changes.

### Conclusions

Hypernatremia can shorten repolarization in both WT and ΔKPQ mouse hearts. We attribute part of this shortening effect to I_to_ enhancement by hypernatremia. Widening the perinexus prolongs APD in genetic mouse and acquired guinea pig sodium channel gain-of-function hearts. However, the combination of hypernatremia and perinexal widening fails to exacerbate APD prolongation in Na_v_GOF mouse hearts due to increased I_to_ in this species. Therefore, the effect of altered extracellular sodium must be considered in studies of repolarization involving sodium channel and transient outward-potassium channel remodeling.

## ACKNOWLEDGEMENT

We would like to thank Dr. Brian P. Delisle at University of Kentucky for generously providing the scn5a-ΔKPQ mice used in this study.

---

**Figure S1.**
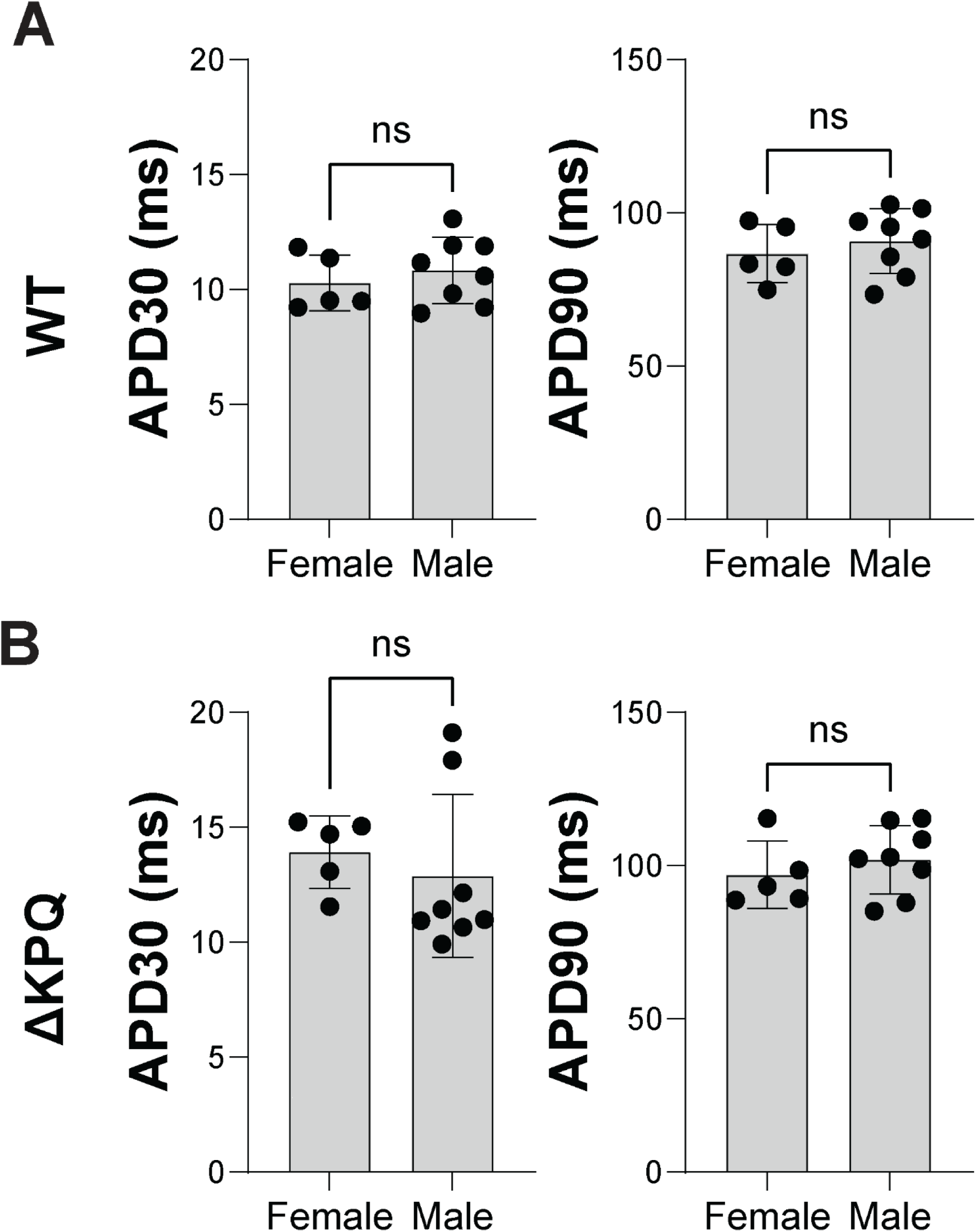
No sex-dependent differences in APD were found in either WT or ΔKPQ hearts. A. APD30 and APD90 from female and male WT hearts (female: n=5; male: n=8). B. APD30 and APD90 from female and male ΔKPQ hearts (female: n=5; male: n=8). *p<0.05 by unpaired Student’s *t* test.

## REFERENCES

1. Chen-Izu Y, Shaw RM, Pitt GS, Yarov-Yarovoy V, Sack JT, Abriel H, Aldrich RW, Belardinelli L, Cannell MB, Catterall WA, et al. Na+ channel function, regulation, structure, trafficking and sequestration. J Physiol. 2015;593:1347–1360. doi: 10.1113/jphysiol.2014.281428

2. Westenbroek RE, Bischoff S, Fu Y, Maier SK, Catterall WA, Scheuer T. Localization of sodium channel subtypes in mouse ventricular myocytes using quantitative immunocytochemistry. J Mol Cell Cardiol. 2013;64:69–78. doi: 10.1016/j.yjmcc.2013.08.004

3. Nuyens D, Stengl M, Dugarmaa S, Rossenbacker T, Compernolle V, Rudy Y, Smits JF, Flameng W, Clancy CE, Moons L, et al. Abrupt rate accelerations or premature beats cause life-threatening arrhythmias in mice with long-QT3 syndrome. Nat Med. 2001;7:1021–1027. doi: 10.1038/nm0901-1021

4. Greer-Short A, George SA, Poelzing S, Weinberg SH. Revealing the Concealed Nature of Long-QT Type 3 Syndrome. Circ Arrhythm Electrophysiol. 2017;10:e004400. doi: 10.1161/CIRCEP.116.004400

5. Nowak MB, Greer-Short A, Wan X, Wu X, Deschenes I, Weinberg SH, Poelzing S. Intercellular Sodium Regulates Repolarization in Cardiac Tissue with Sodium Channel Gain of Function. Biophys J. 2020;118:2829–2843. doi: 10.1016/j.bpj.2020.04.014

6. Wu X, Hoeker GS, Blair GA, King DR, Gourdie RG, Weinberg SH, Poelzing S. Hypernatremia and intercalated disc edema synergistically exacerbate long-QT syndrome type 3 phenotype. Am J Physiol Heart Circ Physiol. 2021;321:H1042–H1055. doi: 10.1152/ajpheart.00366.2021

7. Rhett JM, Ongstad EL, Jourdan J, Gourdie RG. Cx43 associates with Na(v)1.5 in the cardiomyocyte perinexus. J Membr Biol. 2012;245:411–422. doi: 10.1007/s00232-012-9465-z

8. Boutjdir M, Zhang ZH, Huang B, Chen L, Stergiopoulos N, El-Sherif N. Evidence of Na Current Contribution to the Transient Outward Current in Cardiac Ventricular Myocytes. J Cardiovasc Pharmacol Ther. 1996;1:149–158. doi: 10.1177/107424849600100209

9. Zygmunt AC, Goodrow RJ, Antzelevitch C. Sodium effects on 4-aminopyridine-sensitive transient outward current in canine ventricular cells. Am J Physiol. 1997;272:H1–11. doi: 10.1152/ajpheart.1997.272.1.H1

10. Dukes ID, Morad M. The transient K+ current in rat ventricular myocytes: evaluation of its Ca2+ and Na+ dependence. J Physiol. 1991;435:395–420. doi: 10.1113/jphysiol.1991.sp018516

11. Nakayama T, Irisawa H. Transient outward current carried by potassium and sodium in quiescent atrioventricular node cells of rabbits. Circ Res. 1985;57:65–73. doi: 10.1161/01.res.57.1.65

12. Patel SP, Campbell DL. Transient outward potassium current, ‘Ito’, phenotypes in the mammalian left ventricle: underlying molecular, cellular and biophysical mechanisms. J Physiol. 2005;569:7–39. doi: 10.1113/jphysiol.2005.086223

13. Coraboeuf E, Coulombe A, Deroubaix E, Hatem S, Mercadier JJ. [Transient outward potassium current and repolarization of cardiac cells]. Bull Acad Natl Med. 1998;182:325–333; discussion 333-325.

14. Hoppe UC, Johns DC, Marban E, O’Rourke B. Manipulation of cellular excitability by cell fusion: effects of rapid introduction of transient outward K+ current on the guinea pig action potential. Circ Res. 1999;84:964–972. doi: 10.1161/01.res.84.8.964

15. Hume JR, Uehara A. Ionic basis of the different action potential configurations of single guinea-pig atrial and ventricular myocytes. J Physiol. 1985;368:525–544. doi: 10.1113/jphysiol.1985.sp015874

16. Inoue M, Imanaga I. Masking of A-type K+ channel in guinea pig cardiac cells by extracellular Ca2+. Am J Physiol. 1993;264:C1434–1438. doi: 10.1152/ajpcell.1993.264.6.C1434

17. Zicha S, Moss I, Allen B, Varro A, Papp J, Dumaine R, Antzelevich C, Nattel S. Molecular basis of species-specific expression of repolarizing K+ currents in the heart. Am J Physiol Heart Circ Physiol. 2003;285:H1641–1649. doi: 10.1152/ajpheart.00346.2003

18. Vyas S, Saini AG, Kaur A, Singh P, Jayashree M, Sundaram V, Mukhopadhyay K, Singh P. Neuroimaging Spectrum of Severe Hypernatremia in Infants with Neurological Manifestations. Neuropediatrics. 2021;52:316–325. doi: 10.1055/s-0041-1730938

19. Entz M, 2nd, King DR, Poelzing S. Design and validation of a tissue bath 3-D printed with PLA for optically mapping suspended whole heart preparations. Am J Physiol Heart Circ Physiol. 2017;313:H1190–H1198. doi: 10.1152/ajpheart.00150.2017

20. Xu H, Li H, Nerbonne JM. Elimination of the transient outward current and action potential prolongation in mouse atrial myocytes expressing a dominant negative Kv4 alpha subunit. J Physiol. 1999;519 Pt 1:11–21. doi: 10.1111/j.1469-7793.1999.0011o.x

21. Veeraraghavan R, Hoeker GS, Alvarez-Laviada A, Hoagland D, Wan X, King DR, Sanchez-Alonso J, Chen C, Jourdan J, Isom LL, et al. The adhesion function of the sodium channel beta subunit (beta1) contributes to cardiac action potential propagation. Elife. 2018;7. doi: 10.7554/eLife.37610

22. Ly C, Weinberg SH. Automaticity in ventricular myocyte cell pairs with ephaptic and gap junction coupling. Chaos. 2022;32:033123. doi: 10.1063/5.0085291

23. Bondarenko VE, Szigeti GP, Bett GC, Kim SJ, Rasmusson RL. Computer model of action potential of mouse ventricular myocytes. Am J Physiol Heart Circ Physiol. 2004;287:H1378–1403. doi: 10.1152/ajpheart.00185.2003

24. Clancy CE, Rudy Y. Linking a genetic defect to its cellular phenotype in a cardiac arrhythmia. Nature. 1999;400:566–569. doi: 10.1038/23034

25. Trepanier-Boulay V, St-Michel C, Tremblay A, Fiset C. Gender-based differences in cardiac repolarization in mouse ventricle. Circ Res. 2001;89:437–444. doi: 10.1161/hh1701.095644

26. Saito T, Ciobotaru A, Bopassa JC, Toro L, Stefani E, Eghbali M. Estrogen contributes to gender differences in mouse ventricular repolarization. Circ Res. 2009;105:343–352. doi: 10.1161/CIRCRESAHA.108.190041

27. Wu X, King DR, Hoeker GS, Wan X, Deschenes I, Johnstone SR, Gourdie RG, Weinberg SH, Poelzing S. Age-Associated Perinexal Narrowing Masks Consequences of Sodium Channel Gain of Function in Guinea Pig Hearts. JACC Clin Electrophysiol. 2025. doi: 10.1016/j.jacep.2024.12.027

28. Nowak MB, Poelzing S, Weinberg SH. Mechanisms underlying age-associated manifestation of cardiac sodium channel gain-of-function. J Mol Cell Cardiol. 2020;153:60–71. doi: 10.1016/j.yjmcc.2020.12.008

29. Miller JA, Moise N, Weinberg SH. Modeling incomplete penetrance in long QT syndrome type 3 through ion channel heterogeneity: an in silico population study. Am J Physiol Heart Circ Physiol. 2023;324:H179–H197. doi: 10.1152/ajpheart.00430.2022

30. Baartscheer A, Schumacher CA, Coronel R, Fiolet JW. The Driving Force of the Na/Ca-Exchanger during Metabolic Inhibition. Front Physiol. 2011;2:10. doi: 10.3389/fphys.2011.00010

31. Armoundas AA, Hobai IA, Tomaselli GF, Winslow RL, O’Rourke B. Role of sodium-calcium exchanger in modulating the action potential of ventricular myocytes from normal and failing hearts. Circ Res. 2003;93:46–53. doi: 10.1161/01.RES.0000080932.98903.D8

32. Colinas O, Gallego M, Setien R, Lopez-Lopez JR, Perez-Garcia MT, Casis O. Differential modulation of Kv4.2 and Kv4.3 channels by calmodulin-dependent protein kinase II in rat cardiac myocytes. Am J Physiol Heart Circ Physiol. 2006;291:H1978–1987. doi: 10.1152/ajpheart.01373.2005

33. Tessier S, Karczewski P, Krause EG, Pansard Y, Acar C, Lang-Lazdunski M, Mercadier JJ, Hatem SN. Regulation of the transient outward K(+) current by Ca(2+)/calmodulin-dependent protein kinases II in human atrial myocytes. Circ Res. 1999;85:810–819. doi: 10.1161/01.res.85.9.810

34. Deschenes I, Tomaselli GF. Modulation of Kv4.3 current by accessory subunits. FEBS Lett. 2002;528:183–188. doi: 10.1016/s0014-5793(02)03296-9

35. Deschenes I, Armoundas AA, Jones SP, Tomaselli GF. Post-transcriptional gene silencing of KChIP2 and Navbeta1 in neonatal rat cardiac myocytes reveals a functional association between Na and Ito currents. J Mol Cell Cardiol. 2008;45:336–346. doi: 10.1016/j.yjmcc.2008.05.001

36. Wulfers EM, Moss R, Lehrmann H, Arentz T, Westermann D, Seemann G, Odening KE, Steinfurt J. Whole-heart computational modelling provides further mechanistic insights into ST-elevation in Brugada syndrome. Int J Cardiol Heart Vasc. 2024;51:101373. doi: 10.1016/j.ijcha.2024.101373

37. Rosati B, Dong M, Cheng L, Liou SR, Yan Q, Park JY, Shiang E, Sanguinetti M, Wang HS, McKinnon D. Evolution of ventricular myocyte electrophysiology. Physiol Genomics. 2008;35:262–272. doi: 10.1152/physiolgenomics.00159.2007

38. Giudicessi JR, Ye D, Tester DJ, Crotti L, Mugione A, Nesterenko VV, Albertson RM, Antzelevitch C, Schwartz PJ, Ackerman MJ. Transient outward current (I(to)) gain-of-function mutations in the KCND3-encoded Kv4.3 potassium channel and Brugada syndrome. Heart Rhythm. 2011;8:1024–1032. doi: 10.1016/j.hrthm.2011.02.021

39. George SA, Hoeker G, Calhoun PJ, Entz M, 2nd, Raisch TB, King DR, Khan M, Baker C, Gourdie RG, Smyth JW, et al. Modulating cardiac conduction during metabolic ischemia with perfusate sodium and calcium in guinea pig hearts. Am J Physiol Heart Circ Physiol. 2019;316:H849–H861. doi: 10.1152/ajpheart.00083.2018

40. Adams WP, Raisch TB, Zhao Y, Davalos R, Barrett S, King DR, Bain CB, Colucci-Chang K, Blair GA, Hanlon A, et al. Extracellular Perinexal Separation Is a Principal Determinant of Cardiac Conduction. Circ Res. 2023;133:658–673. doi: 10.1161/CIRCRESAHA.123.322567

41. Blair GA, Wu X, Bain C, Warren M, Hoeker GS, Poelzing S. Mannitol and hyponatremia regulate cardiac ventricular conduction in the context of sodium channel loss of function. Am J Physiol Heart Circ Physiol. 2024;326:H724–H734. doi: 10.1152/ajpheart.00211.2023

